# Early transcriptional responses after dengue vaccination mirror the response to natural infection and predict neutralizing antibody titers

**DOI:** 10.1101/314377

**Authors:** Stephen J. Popper, Fiona R. Strouts, Janet C. Lindow, Henry K. Cheng, Magelda Montoya, Angel Balmaseda, Anna P. Durbin, Stephen S. Whitehead, Eva Harris, Beth D. Kirkpatrick, David A. Relman

## Abstract

**Background:** Several promising live attenuated virus (LAV) dengue vaccines are in development, but information about innate immune responses and early correlates of protection are lacking.

**Methods:** We characterized human genome-wide transcripts in whole blood from 10 volunteers at 11 time-points after immunization with the dengue virus type 3 (DENV-3) component of the NIH dengue vaccine candidate TV003 and from 30 hospitalized children with acute primary DENV-3 infection. We compared day-specific gene expression patterns with subsequent neutralizing antibody (NAb) titers.

**Results:** The transcriptional response to vaccination was largely confined to days 5–20 and was dominated by an interferon-associated signature and a cell cycle signature that peaked on days 8 and 14, respectively. Changes in transcript abundance were much greater in magnitude and scope in symptomatic natural infection than following vaccination (maximum fold-change >200 versus 21 post-vaccination; 3,210 versus 286 transcripts with significant fold-change), but shared gene modules were induced in the same sequence. The abundance of 131 transcripts on days 8 and 9 post-vaccination was strongly correlated with NAb titers measured 6 weeks post-vaccination.

**Conclusions:** LAV dengue vaccination elicits early transcriptional responses that mirror those found in symptomatic natural infection and provide candidate early markers of protection against DENV infection.

Clinical Trial Registration Number: NCT00831012 (available at clinicaltrials.gov)

Summary: Interferon- and cell cycle-associated gene transcript abundance levels in the peripheral blood of dengue vaccine recipients on days 8 and 9 post-vaccination were associated with dengue neutralizing antibody titers on day 42, and mirrored responses in primary dengue infection, suggesting the possibility of predicting protective immunity.

## Background

Each year, the four dengue virus serotypes (DENV-1–4) infect an estimated 390 million individuals globally [1]. While most of these infections are asymptomatic, approximately 100 million individuals develop clinically apparent disease, from uncomplicated fever to life-threatening illness. Despite the high disease burden, there are no licensed therapeutics for DENV infection. Several promising candidate dengue vaccines are in Phase III clinical trials, and the live attenuated chimeric dengue vaccine Dengvaxia^™^ was recently licensed for use in children 9 years of age and older in DENV endemic areas. However, the efficacy and duration of protection were limited or uncertain, and DENV-naïve vaccine recipients were hospitalized for dengue and severe dengue at a higher rate than placebo recipients, possibly due to antibody-dependent enhancement (ADE) [2].

Studies of natural DENV infection and flavivirus LAVs have identified immune responses needed for protection against dengue disease. Pre-existing neutralizing antibody (NAb) titers correlate with a lack of symptomatic disease in subsequent infections [3–6] and are used as the primary measure of candidate vaccine immunogenicity. However, the risk of severe disease is elevated after a second infection with a heterotypic dengue virus [7]. The recognition of effective homotypic immunity after natural infection has led to a common vaccine development strategy of inducing homotypic NAbs to all four serotypes simultaneously.

Little is known about the role of early innate immune responses in enhancing NAb production and promoting protective immune memory against dengue. Studies of innate immunity have been hampered by the difficulty in identifying individuals with early infection, when innate immune responses are most active, particularly those with mild or subclinical infections. Trials of LAVs provide a unique opportunity to examine early immune responses in a setting where the time, dose, and viral serotype are known. Genome-wide transcript responses to vaccines have provided important clues about early steps in the generation of humoral and cellular immunity [8–13]. Transcript profiling of peripheral blood also incorporates information from cell populations that are difficult to examine in clinical settings, and has led to signatures associated with dengue disease severity, identified links between innate responses and humoral immunity in secondary DENV infection, and illustrated the dynamic nature of these responses [14–20].

In this study, we characterized the transcript response to rDEN3Δ30/31, the DENV-3 component of TV003, a tetravalent live attenuated vaccine candidate developed by NIH. TV003 is a single-dose vaccine that has proven to be both safe and immunogenic and is being evaluated in a Phase III efficacy trial [21,22]. We examined temporal changes in transcript abundance and identified early signatures correlated with NAb titers measured six weeks post-vaccination. We also compared these results with transcript patterns we observed in patients with symptomatic wild-type primary DENV-3 infection. Despite the anticipated differences in the magnitude of expression, we observed the induction of common gene expression programs in the same temporal sequence, with a similar relationship to the induction of NAb. These results reveal candidate biomarkers of early protective DENV immune responses against dengue and suggest a path towards validation and deployment.

## Methods

### Vaccine study population

Samples for this study were collected from a Phase I clinical trial of the live attenuated dengue vaccine rDEN3Δ30/31–7164 (DENV-3), described previously [23]. Briefly, healthy, flavivirus-naïve adult volunteers were randomized to receive a single 0.5 ml subcutaneous dose of 1,000 PFU of DENV-3 vaccine or a placebo (0.5 ml of vaccine diluent). Blood samples including whole blood for RNA profiling were collected immediately prior to vaccination and on days 2, 5, 6, 8, 9, 12,14, 20, 29, 42 and 180. Samples from each of these time-points were available from nine of ten vaccinees and from all placebo recipients. Subject 9 had samples available for all days except days 8 and 12; 166 samples in total were used for analysis. Serum virus titers (viremia) were measured using a standard plaque assay as described previously [24]. Serum NAb titer was determined by 60% plaque reduction (PRNT_60_) [25]. Seroconversion was defined by a ≥4-fold increase in PRNT_60_ on study day 28 or 42 relative to day 0 and corresponds to a post-vaccination titer >10 [23].

### Dengue patient population

Patients (6 months to 14 years old) presenting with fever and suspected dengue during the 2010 dengue season were enrolled at the Hospital Infantil Manuel de Jesús Rivera in Managua, Nicaragua. Inclusion criteria, recruitment, and laboratory testing have been described previously [26] (see Supplementary Information). Blood samples from healthy subjects were collected as part of a separate prospective cohort study in which healthy children in the same general population were enrolled without regard to dengue status [27].

### Ethics statement

The trial of rDEN3Δ30/31 was approved by the Committee for Human Research at the University of Vermont, and written informed consent was obtained from all subjects following a review of risks and benefits and a comprehension assessment. The study in Nicaragua was approved by the Institutional Review Boards of the University of California, Berkeley, and the Nicaraguan Ministry of Health, and by the Stanford University Administrative Panel on Human Subjects in Medical Research. For further details, see Supplementary Information.

### RNA sample processing and transcriptome analysis

PAXgene RNA was amplified and hybridized to Human Exonic Evidence Based Oligonucleotide microarrays [14]. Microarray data were submitted to the Princeton University MicroArray database for normalization and gene filtering and are deposited at Gene Expression Omnibus (http://www.ncbi.nlm.nih.gov/geo/; accession numbers GSE96656 and GSE98053). A full description of both sample processing and analysis steps is available in the Supplementary Information.

## Results

### Temporal patterns of the transcriptional responses to live dengue vaccination

To identify the temporal pattern of the early human transcriptional response to dengue vaccination, we examined changes in genome-wide transcript abundance in serial whole blood samples from 10 volunteers infected with 1,000 plaque forming units (pfu) of rDEN3Δ30/31, the dose included in TV003, and four volunteers inoculated with placebo (L-15 medium). Nine of ten vaccinees seroconverted 28 days post-vaccination, defined as a 60% plaque reduction neutralization titer (PRNT_60_) >10 (Table 1). Four of the vaccinees had low-level viremia on one or more days within the first 10 days post-vaccination, five developed a mild maculopapular rash, and none were febrile. The four placebo recipients remained seronegative for DENV serotypes.

**Table 1.**
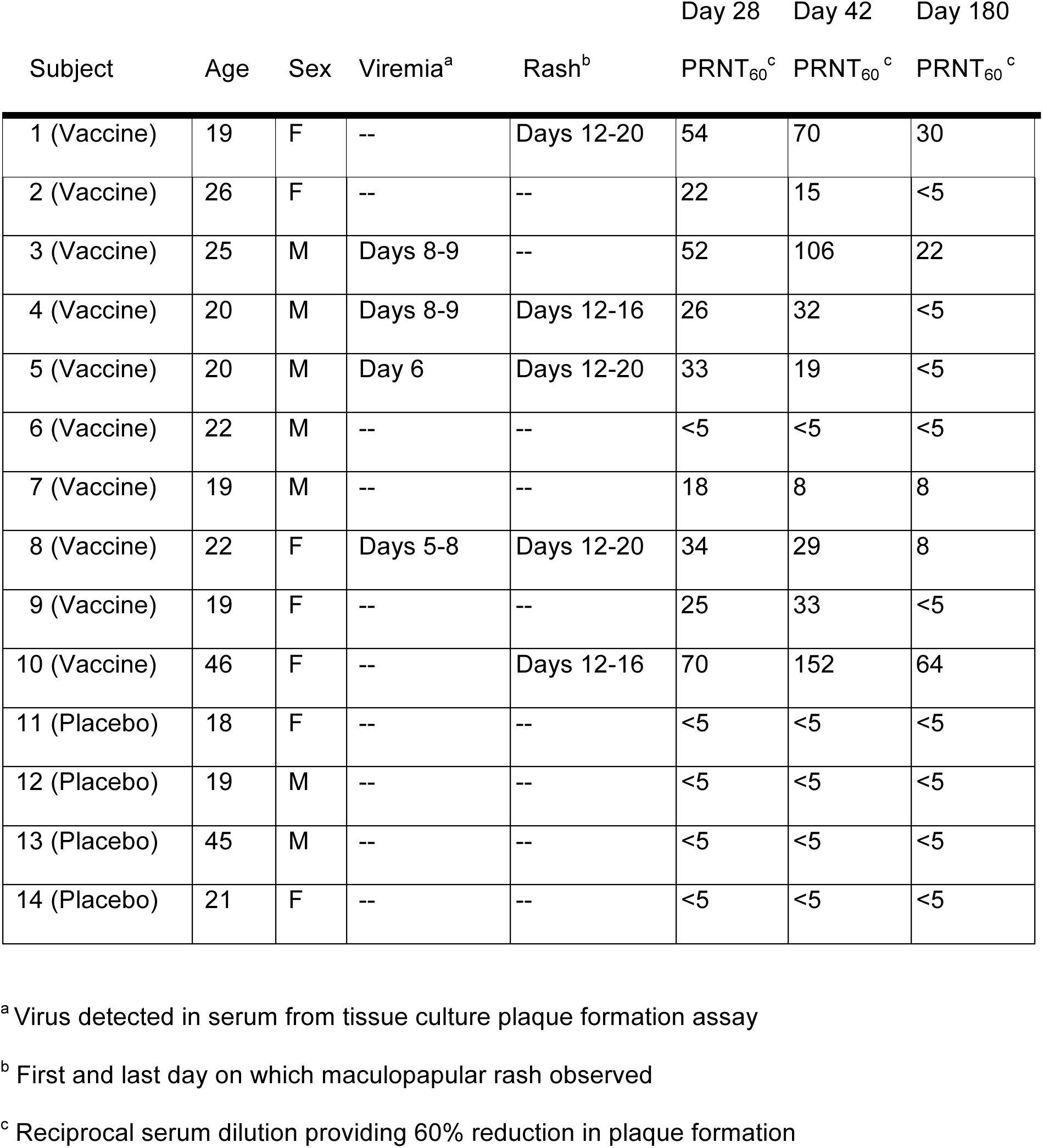
Characteristics of subjects in vaccine trial

We collected whole blood for isolation of RNA immediately before vaccination (day 0), and on days 2, 5, 6, 8, 9, 12, 14, 20, 29, 42 and 180 post-vaccination from all volunteers and measured genome-wide transcript abundance levels. Data were available for eight of the nine participants who seroconverted. For each of these eight subjects, we compared transcript abundances for each post-vaccination day with those for the matched pre-vaccination sample (see Supplementary Information). Almost all significant changes in transcript abundance occurred 5–20 days after vaccination, with a peak of 161 and 156 transcripts changing in abundance (days 8 and 9, respectively), and 286 transcripts with a significant change in abundance on at least one day (Figure 1). Fewer transcripts met criteria for significance when comparing vaccinees to placebo recipients (n=131), but the direction of change for 271 of the 286 transcripts from vaccinees was the same whether the comparison was with day-matched placebo recipients or each subject’s baseline sample (Supplementary Figure 1).

**Figure 1.**
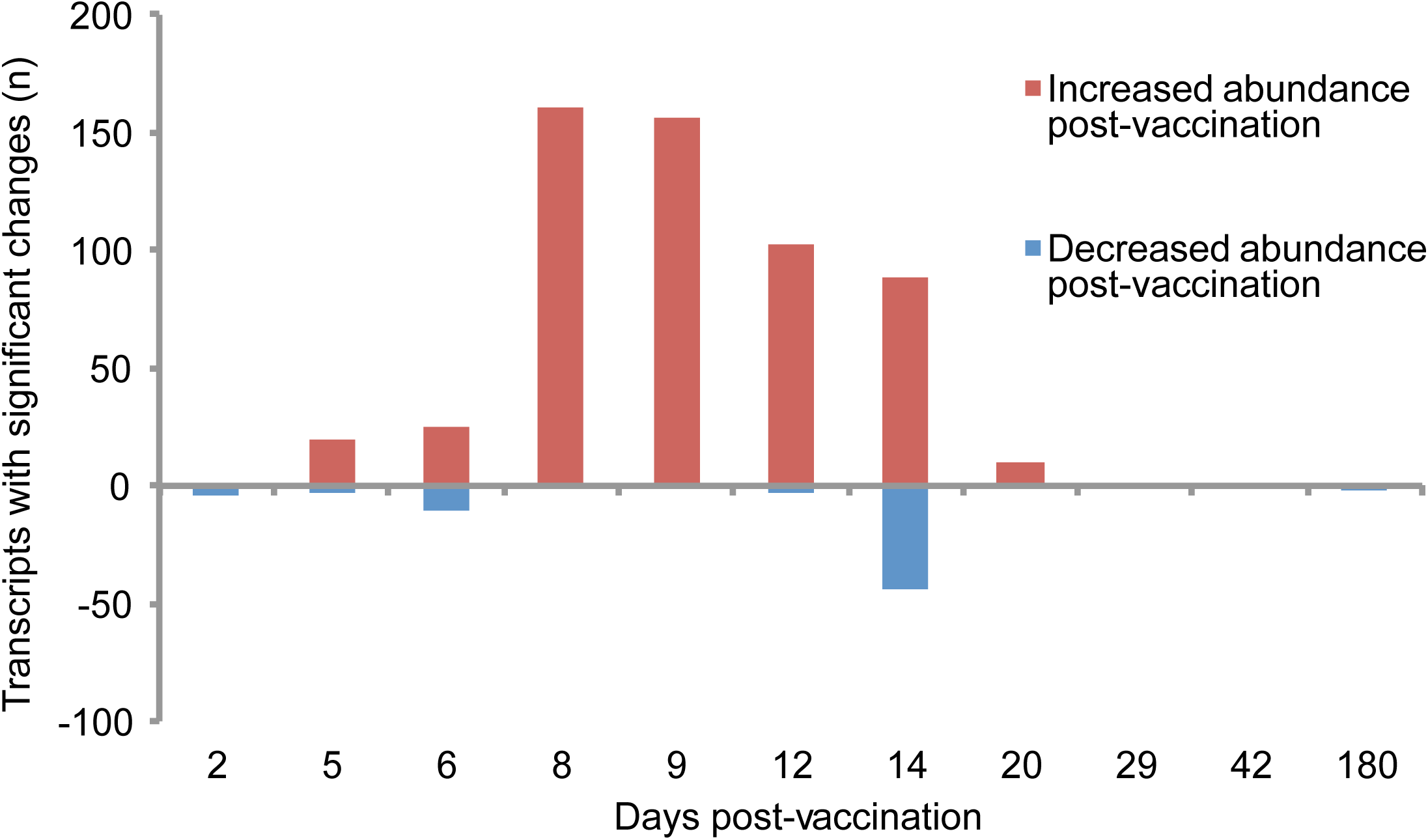
Significant differences in transcript abundance post-vaccination (FDR<1%; minimum 2-fold change compared to pre-vaccination sample).

To infer the functional implications of these changes in transcript abundance, we used hierarchical clustering to organize the transcripts and compared gene membership in Gene Ontology and the KEGG pathways using the DAVID bioinformatics resource [28]. Gene transcripts were grouped in three clusters (Figure 2 and Supplementary Figure 2). Transcripts in Cluster 1 were more abundant after vaccination (Figure 2C), peaked on days 8 and 9 post vaccination, and included canonical interferon-stimulated gene (ISG) transcripts; IFI44, IFI44 L, IFI27, HERC5, IFIT1, USP18, and ISG15 transcripts all increased 10- to 22-fold compared to baseline. Cluster 1 was strongly enriched for genes involved in the innate immune response to viruses and highly enriched for genes we previously found to be expressed after treatment of PBMCs with type I interferon (p<1E-36) [29].

**Figure 2.**
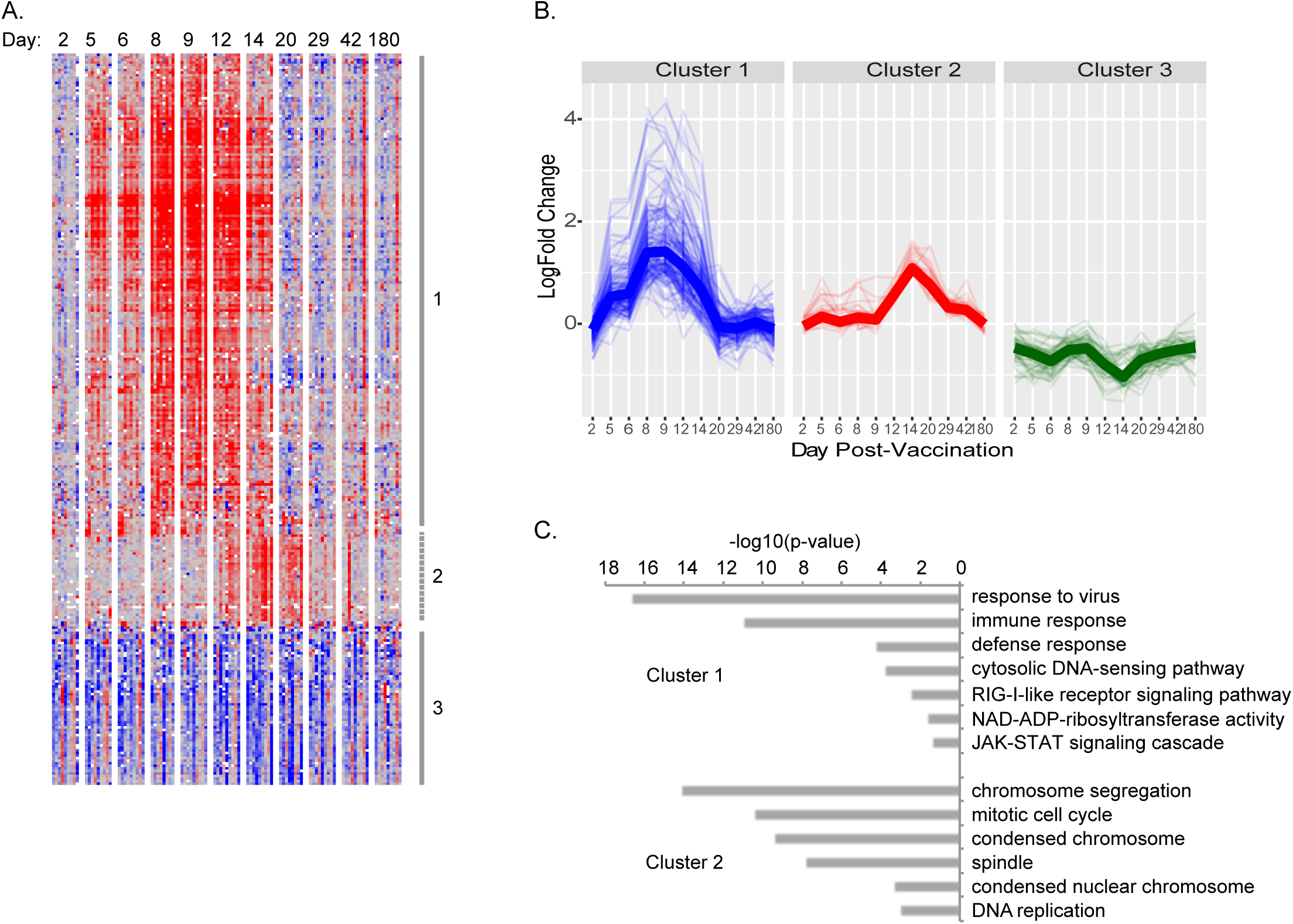
Changes in transcript abundances over time in vaccinees. A) Hierarchical clustering of the 286 transcripts whose abundance was significantly different from baseline on more than one day. Lines and numbers to the right of the heatmap mark sets of co-expressed genes (average cluster r>0.5). B) Change over time in abundance for each transcript in each gene cluster. Heavy line indicates median expression of all genes in each cluster. C) Gene ontologies associated with gene clusters described in (A) and (B). There were no significant gene ontologies for Cluster 3.

Gene transcripts in Clusters 2 and 3 showed maximal changes on day 14, with Cluster 2 transcripts increasing and Cluster 3 transcripts decreasing in abundance from baseline (Figure 2A and 2B). Cluster 2 included TYMS, CEP55, CCNA2, and NEK2, whose genes products are involved in DNA replication and cell division, and other genes associated with mitosis (p<2E-9, Figure 2C). Genes in Cluster 3 were enriched in both reticulocytes (p=1E-20) and neutrophils (p=2E-7) [30]. We did not measure reticulocyte counts, but we did measure neutrophils and the relative neutrophil abundance in vaccinees did not change significantly with time (p=0.55, paired t-test), suggesting that decreased expression of these genes was not due to decreased neutrophil abundance.

### Changes observed after vaccination are a subset of those observed in natural symptomatic DENV-3 infection

To establish which features of the early response to vaccination are shared with the response to natural symptomatic infection, we examined transcript responses in Nicaraguan children hospitalized with acute dengue. We previously demonstrated that a history of previous DENV exposure is the most prominent source of variation in gene expression in dengue patients [14]. To ensure that DENV immune status, as well as serotype, did not confound our analysis, we identified 30 children diagnosed with acute primary DENV-3 infection during a single year (24 with dengue fever and 6 with dengue hemorrhagic fever; Supplementary Table 1), and compared transcript abundance in whole blood with measurements from 9 healthy individuals. Principal components analysis confirmed previous findings that there are significant day-to-day changes in the transcript response to natural infection [14,31] (Supplementary Figure 4); thus, we subsequently performed analyses stratified by day of fever. There were no significant differences in transcript abundance between children with dengue fever and children with dengue hemorrhagic fever after matching for sex and day of fever, which is generally consistent with previous studies in the same population [14,18], although the small sample size could affect the results of our analysis.

Despite having fewer days available for comparison and lacking baseline samples for each patient, we identified many more transcripts with significant changes in abundance post-infection compared to those found in vaccinees: among the 20,623 transcripts measured in both datasets, we identified 3,210 transcripts that differed significantly on at least one day of fever, compared with 278 transcripts following vaccination (Figure 3A, Supplementary Figure 5A). The magnitude of the maximum change in abundance post-infection was also nearly 10-fold greater: there was a 200-fold difference post-infection compared to a maximum 21-fold difference post-vaccination (Figure 3B). The transcripts with the greatest differences in relative abundance during natural infection were MT2A (242-fold) and USP18 (183-fold), both of which are interferon-induced; HESX1 (150-fold), which is expressed in activated dendritic cells; and SPAT2SL (137-fold), which may be involved in activation and differentiation of multiple cell types.

**Figure 3.**
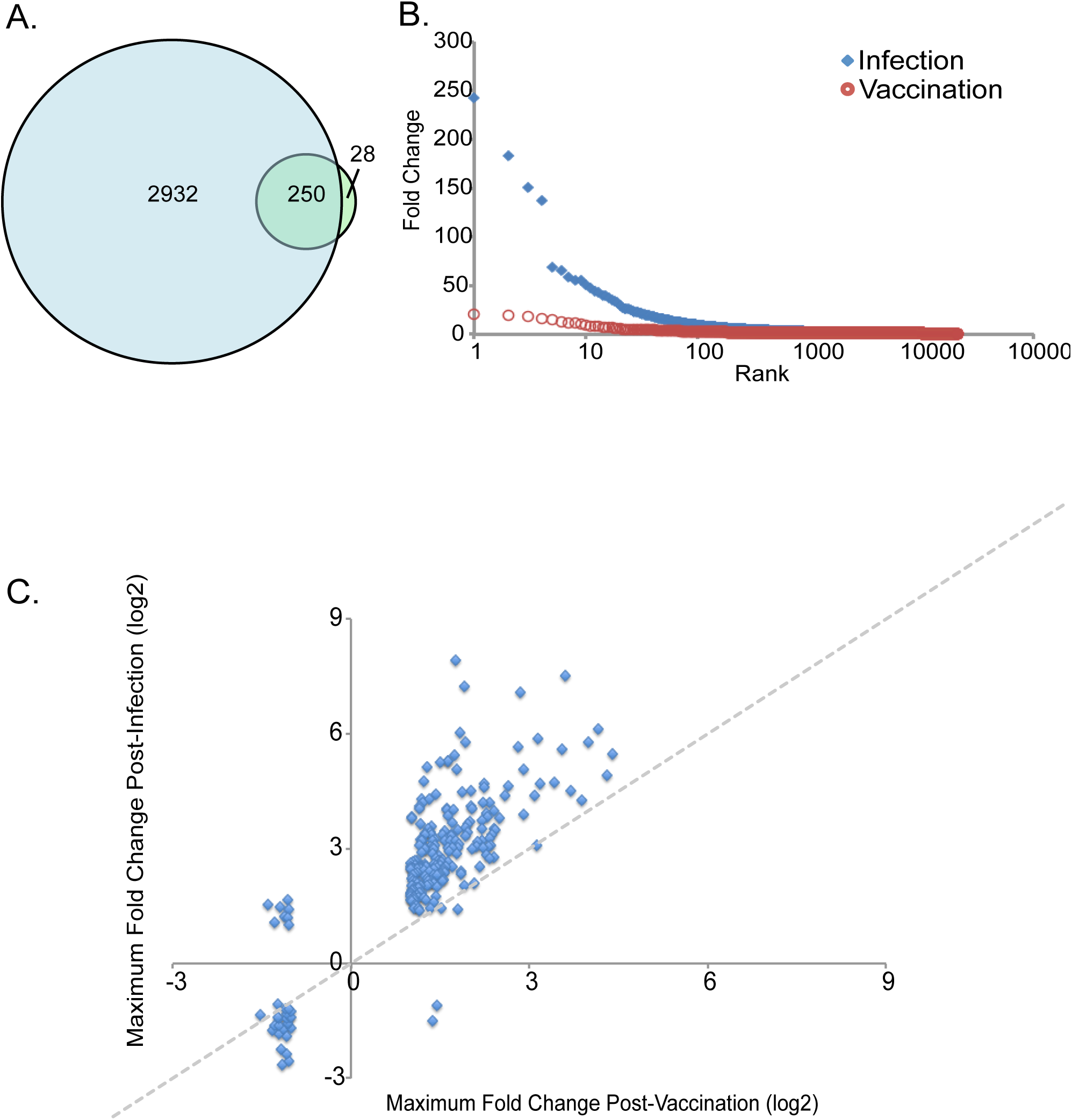
Comparison of post-vaccination and post-infection transcript abundance changes. A) Transcripts with significant changes on days 2, 3, 4, or 5 of fever in patients with primary DENV-3 infection (blue circle) and on any day post-vaccination (green circle). Numbers indicate transcripts unique to vaccination, infection, or shared (overlap, n=246). B) Maximum fold-change in transcript abundance following vaccination (red circles) or during infection (blue diamonds). C) Maximum fold-change in abundance for transcripts with significant changes post-vaccination or during infection. Dotted diagonal line at equal fold change included for reference.

Despite differences in response magnitude and number, the response following natural symptomatic infection included 90% (250/278) of transcripts that changed after vaccination, and the direction of change was the same for 96% of these transcripts (240/250) (Figure 3C). The transcripts that changed the most post-vaccination (IFI44, IFI44 L, IFI27 and HERC5) were among the 20 transcripts with the biggest differences in abundance following natural infection, and relative increases in transcript abundance were strongly correlated across the two groups (Spearman r^2^ = 0.75).

### Responses to dengue vaccination and symptomatic natural infection share a common temporal sequence

We used gene set enrichment analysis and information from all measured transcripts to identify 141 blood transcript gene modules that changed in abundance following either immunization or infection [8] (FDR<1%). Many of these modules demonstrated similar changes in both vaccinees and patients (Figure 4A). Modules enriched for ISG expression were elevated on days 5–14 post-vaccination and were also persistently elevated after natural DENV infection. Modules representing monocyte-associated transcripts were elevated on days 1–3 of natural infection and on days 8–9 post-vaccination, while modules associated with the mitotic cell cycle were elevated on later days in both groups, with the highest levels on day 5 of natural infection and on day 14 post-vaccination. When we compared the overall profiles of the gene modules in the two groups, we found that the responses to natural infection on fever days 1–3 were most similar to responses to vaccination on days 8–9 (Pearson’s r ≥0.60; peak on day 9), while fever day 4 was most similar to vaccination day 12 (r>0.75, peak on day 12), and fever day 5 was most similar to vaccination day 14 and subsequent time-points (r≥0.70, peak on 14) (Figure 4B, Supplementary Dataset 1). Thus, the enrichment of common modules in the same sequence indicates a similar progression in the early host response to vaccination and to natural infection.

**Figure 4.**
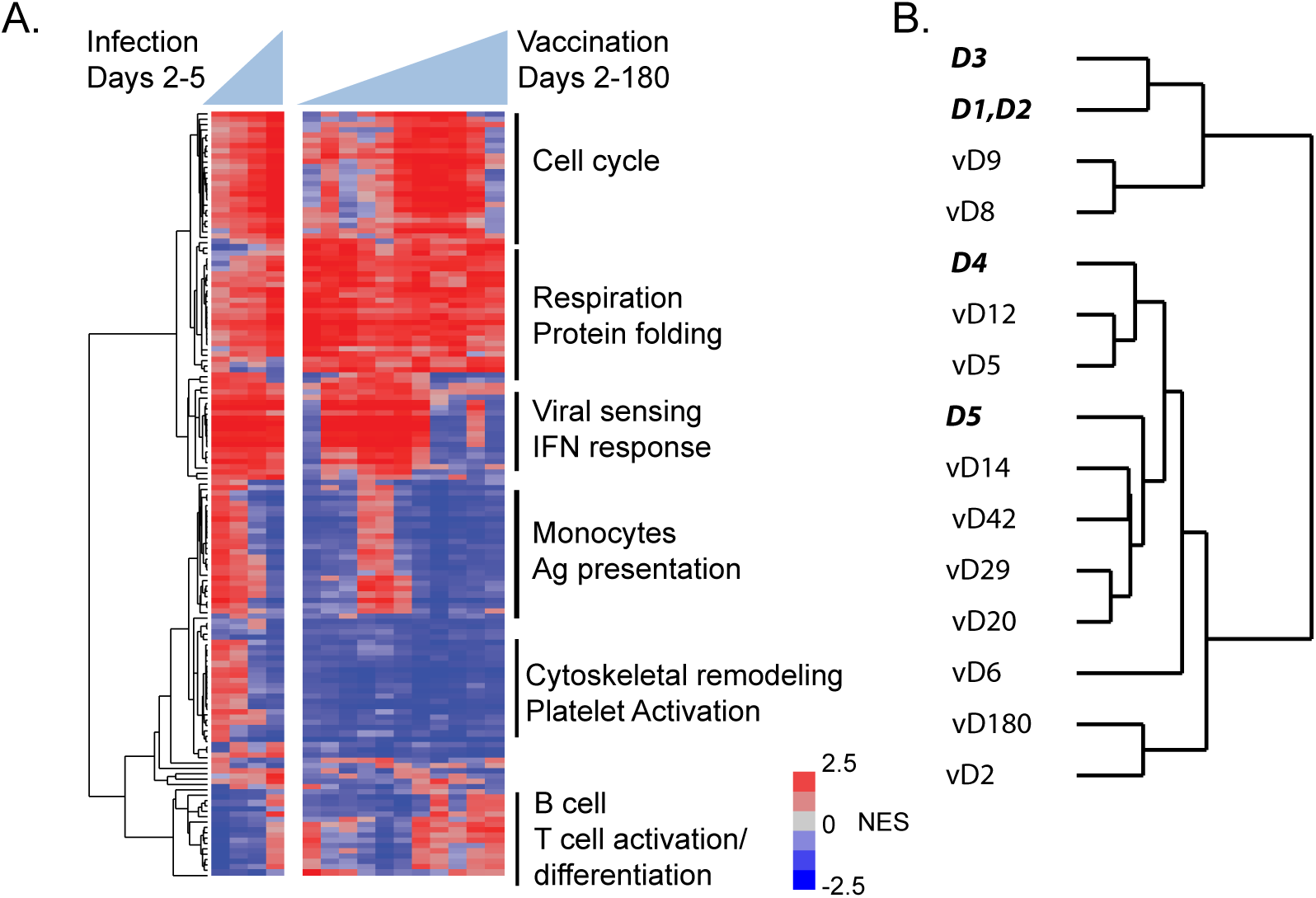
Gene modules affected by DENV vaccination and natural infection. A) Blood transcript modules with transcripts that were significantly up- or down-regulated on at least one day (FDR <1%) were hierarchically clustered. NES; normalized enrichment score. Vertical lines on right denote module clusters described in the text. B) Hierarchical clustering of each day post-vaccination or post-infection using the NES from (A). Days in bold italics represent days of fever for infected patients; days preceded by “v” represent days post-vaccination.

We note there was also a cluster of 16 gene modules, six associated with platelet activation and cytoskeletal remodeling, that were elevated in natural infection but not vaccinees (Figure 4A, Supplementary Dataset 1). Previous studies have demonstrated that platelet activation and TGFβ expression are elevated in DENV infection and higher in patients with more severe disease [32]. TGFβ, which is expressed at high levels in platelets [33], was elevated on fever days 1–2 in dengue patients but was never elevated post-vaccination (Supplementary Figure 6).

### Early transcriptional responses linked to neutralizing antibody production

DENV-specific NAbs are the primary endpoint for assessing vaccine responses in clinical trials and are associated with protection from both symptomatic infection and severe disease [3–5]. To determine whether changes in host transcript patterns predicted differences in NAb titer we calculated the correlation between the change in abundance of each transcript on each day and the NAb titer on post-vaccination day 42, when NAbs are generally at peak titer (Table 1, Supplementary Figure 7A). During the first 6 days post-vaccination, we found no significant correlations with NAb titer, but by day 8, expression of the ISGs in Cluster 1 positively correlated with the day 42 NAb titer (p<0.01; Figure 5). This correlation was equally strong on day 9, and 131 transcripts were significantly correlated with day 42 NAb titer on both days. Among the individual ISG transcripts most strongly correlated with day 42 NAb titer on both days 8 and 9 was IFI44, the transcript whose abundance changed the most post-vaccination (Supplementary Figure 7B). IFI44 was also elevated at one time-point in each of two placebo recipients, but the timing of elevated expression was different and correlated with unrelated respiratory viral infections in each instance (Supplementary Figure 8). Twelve of the 131 transcripts were also associated with subsequent development of a rash, which was the only significant correlate with positive NAb titer in a clinical trial of TV003 [21] (Supplementary Figure 9). Interestingly, the one vaccinee who failed to develop neutralizing antibodies showed little evidence of increased abundance in Cluster 1 genes (Supplementary Figure 3). The association of interferon-related transcript abundance and later NAb titer diminished on days 12 and 14, but BUB1 (r=0.86) and other transcripts associated with the mitotic cell cycle were correlated with subsequent NAb titers on day 14 (Figure 5, Supplementary Figure 7C).

**Figure 5.**
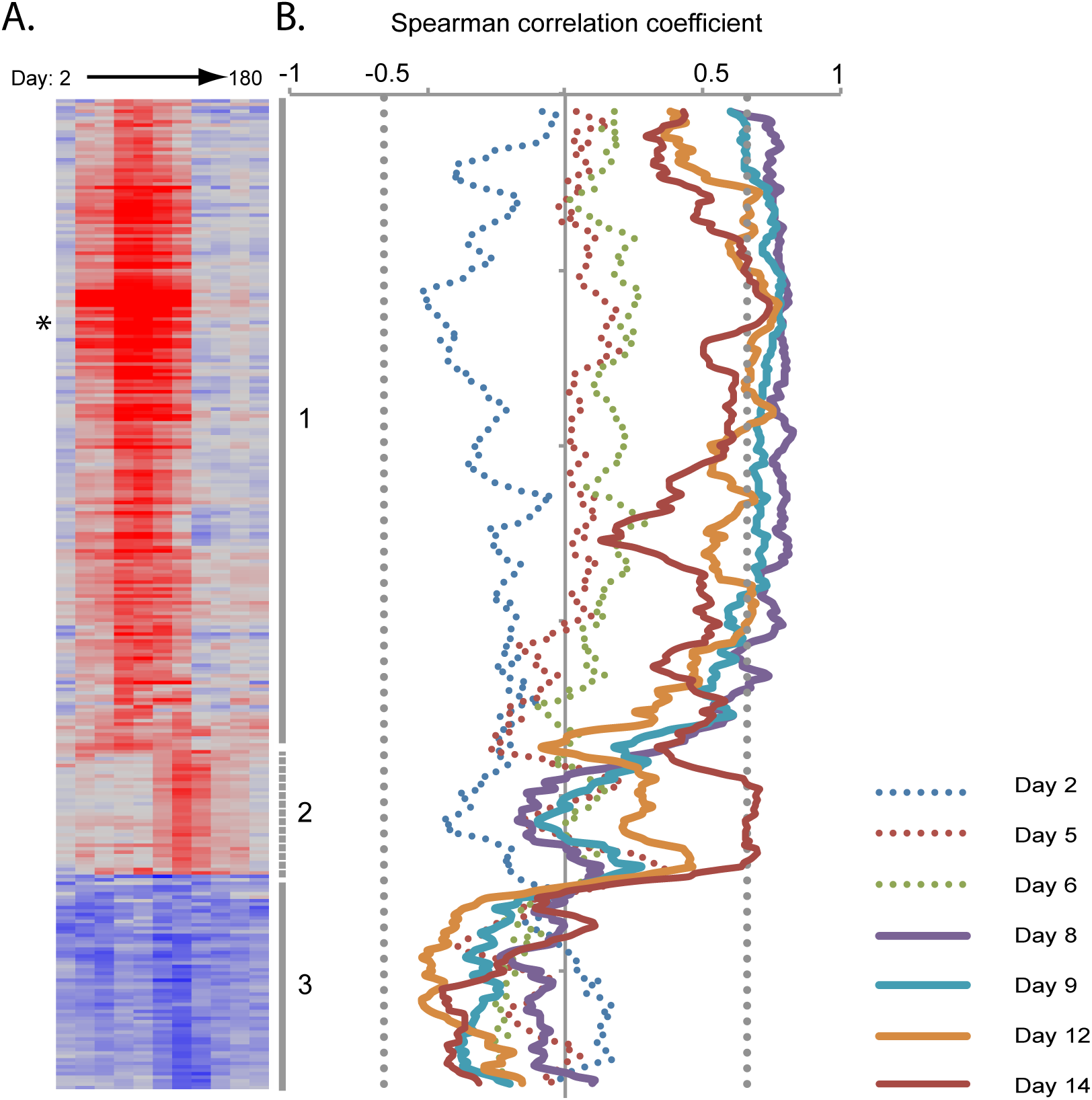
Correlation of transcript abundance and day 42 PRNT_60_ among vaccine recipients. A) Average fold change in abundance by day for all transcripts with significant differences from baseline post-vaccination. Transcripts are ordered and clusters labeled as in Figure 2. Asterisk marks IFI44. B) Spearman correlation of each transcript and day 42 PRNT_60_ using a moving average of window size 9. Solid lines indicate days post-vaccination on which a significant correlation was identified (p<0.01, indicated by vertical dotted grey line).

When we performed similar comparisons for naturally infected patients, we found no transcript clusters significantly correlated with either convalescent or three month NAb titer (Supplementary Figure 5B and 5C). However, the pattern of blood transcript module enrichment indicated a similar relationship between day-specific gene expression and later production of NAb; gene enrichment for both interferon-stimulated and cell cycle-associated gene modules was associated with higher NAb titer in both vaccinees and patients (Figure 6), albeit more weakly in patients, and cell cycle-associated modules were correlated with NAb titer later in both groups.

**Figure 6.**
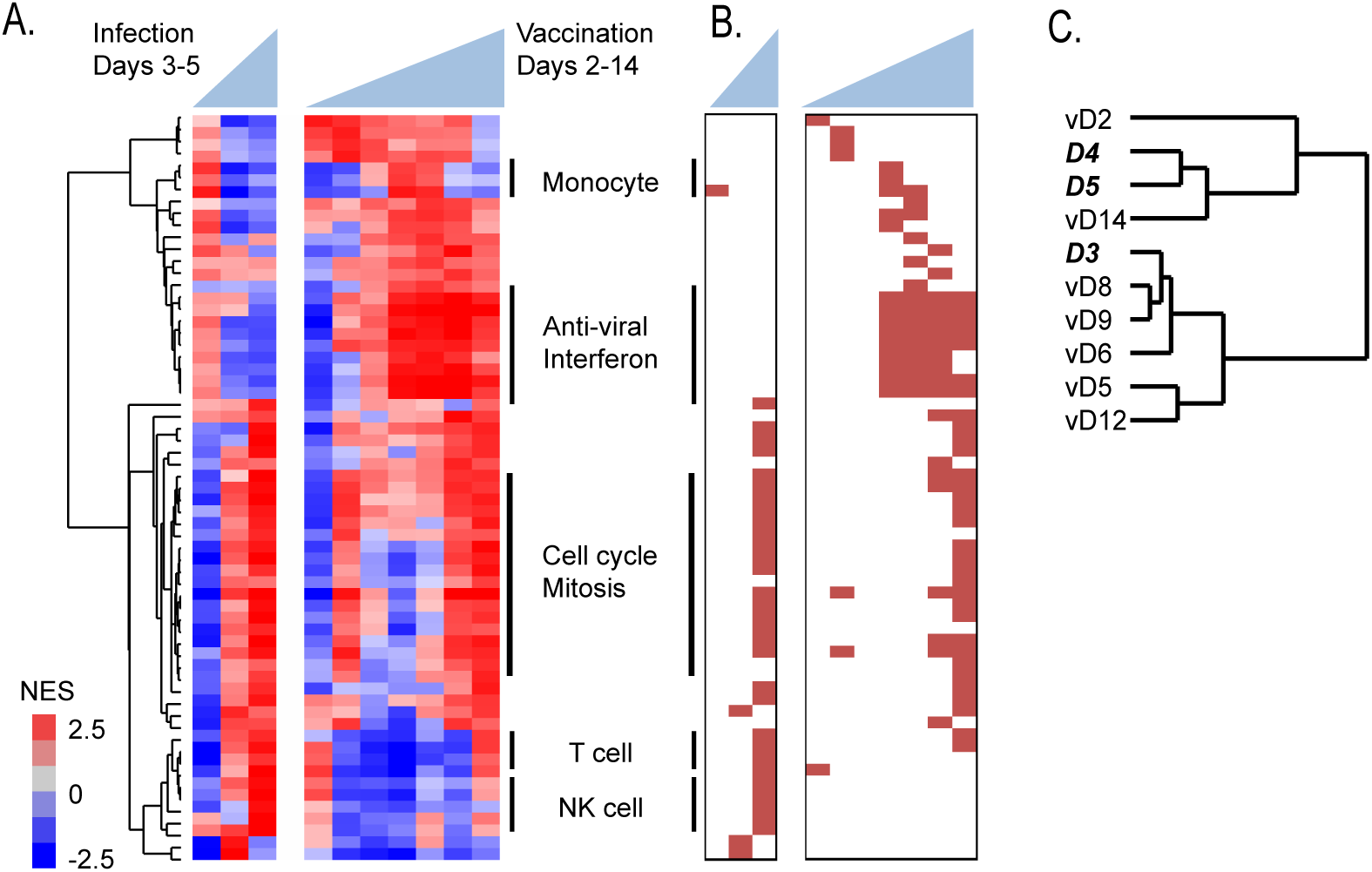
Gene modules correlated with subsequent neutralizing antibody response. A) Blood Transcript Modules that were significantly enriched for transcripts positively correlated with day 42 PRNT_60_ (vaccinees) or convalescent NT_50_ (patients) on at least one day (FDR<1%) were hierarchically clustered. NES; normalized enrichment score. Vertical lines delineate module clusters described in the text. B) Significant modules (FDR<1%) are marked in red. Modules and samples are organized as in (A). C) Hierarchical clustering of gene module expression from each day post-vaccination or post-infection using the NES from (A). Day labels in bold italics represent fever day for infected patients; day labels preceded by “v” represent day post-vaccination.

There are at least three subpopulations of monocytes with distinct transcript profiles [34]; Kwissa et al. identified an increase in CD14^+^CD16^+^ intermediate-phenotype population after secondary DENV infection, and showed that in vitro these cells stimulated formation of the plasmablasts that secrete antibodies weeks after infection, mediated in part by secretion of the ISG cytokine BAFF [19]. In our study, gene set enrichment analysis indicated enrichment of transcripts for both intermediate and nonclassical monocytes at multiple time-points in both vaccinees and patients, while BAFF transcripts were most abundant on fever days 1 and 2 in the patients and days 8 and 9 in the vaccinees (Supplementary Figure 10).

## Discussion

In this study, we used intensive longitudinal sampling to characterize the transcriptional response to dengue vaccination, compared results with those from natural infection with the same DENV serotype, and identified early features that may predict a protective immune response. We found that vaccination and natural infection induced common gene expression programs, and the abundance of individual interferon-stimulated transcripts 8 days post-vaccination was correlated with NAb titers measured five weeks later, representing the earliest identified correlates of a protective adaptive immune response following dengue vaccination.

An interferon response signature has been observed in other studies profiling viral vaccine transcriptional responses. Inactivated influenza and meningococcal vaccines both induce a mild interferon response during the first week post-vaccination, but the response is particularly strong after vaccination with live attenuated vaccines [9,12,35]. We reported that ISG expression was much stronger in cynomolgous macaques infected with wild-type DENV compared to live attenuated virus [35]. Here, we found that ISG expression was much stronger in symptomatic dengue patients than vaccinees, presumably due to higher viral load after infection with wild-type virus. Expression of ISGs was correlated with viral load in the patients, as seen in other studies [19,36]. However, this association did not persist when patients were stratified by day of fever, highlighting the importance of temporal variation in the innate immune response and in viral load, and suggesting that factors in addition to viral replication influence ISG expression. Several studies have found stable inter-individual differences in the response to interferon, suggesting that genetic and environmental features may affect the relationship between viral infection and the interferon response [37,38]. We were not able to assess directly the impact of these features on transcriptional responses in this study, but their contributions to the differences between children with symptomatic infection and vaccinated adults are likely to be much less than the contribution of differences in virulence of vaccine and wild type viruses (see Supplementary Information).

The links between type I interferon production and NAb production probably involve multiple cell types. Plasmacytoid dendritic cells (pDCs) contribute to B cell differentiation and antibody production after viral infection [39]. In this study, increases in monocyte-associated gene expression coincided with ISG expression, and we found features related to multiple monocyte phenotypes in both natural infection and vaccination. Gene module analysis also suggested that T cells were responsible for the increase in cell cycle-associated transcripts two weeks after vaccination that was linked to NAb titers. Future targeted studies of pDCs, monocytes, and T cell populations during the first two weeks post-vaccination will help clarify their role in establishing long-lasting antibody responses. In addition, the link between an early interferon response and later NAb titer was only apparent in natural infection when we used a module analysis approach. This may indicate a plateau, or saturation effect, in the relationship between ISG expression and antibody titer. Alternatively, it may reflect the variability in pathogen dose, prior health status and/or days of infection absent in clinical trials but inherent in observational studies.

Comparison with LAV vaccination also provides a framework for identification of features associated with pathogenic versus non-pathogenic infection. A recent study compared PBMC gene expression in asymptomatic and clinically significant secondary DENV infection and identified differences in antigen presentation and lymphocyte activation [36]. In this study examining whole blood gene expression during primary infection, we found an increased abundance of transcripts associated with platelet activation in natural (pathogenic) infection but not vaccination (non-pathogenic infection), consistent with the hypothesis that platelet activation contributes to dengue pathogenesis [40].

Neutralizing antibody titers were used as an endpoint for these vaccine studies because many studies have shown that these antibodies play an important role in protective immunity. However, recent work has demonstrated that NAbs measured in vitro are an imperfect correlate of in vivo protection [37,38]. Immunity mediated by NAbs may be neither life-long nor sterilizing [43,44] and will be affected by the quality as well as the quantity of NAbs [5,26,45]. Recent studies also highlight a likely role for cytotoxic T cells in mediating protection against DENV reinfection and severe disease [46–49]. The NIH tetravalent vaccine, of which rDEN3Δ30/31 is a component, elicits CD4^+^ T cell responses similar to those seen in natural infection [50]. It will be important to establish whether the early transcript-based features we measured in this study are associated with DENV-specific responses in memory T cell populations.

Our findings reflect the integration of data across multiple time-points and thousands of transcripts, and provide a robust basis for further investigation. We previously found that early interferon-associated transcriptional responses were associated with antibody formation in non-human primates exposed to a tetravalent dengue vaccine [35]. We believe it is likely that the same relationship will exist in humans immunized with tetravalent LAV dengue vaccines. The initiation of Phase 3 clinical trials of TetraVax-DV-TV003 provides an opportunity to validate the relationship between these early responses and NAb titers, and identify specific transcripts as early surrogate markers of both immunogenicity and protection.

## Acknowledgements

We thank Cassandra Ventrone for sample collection and logistics, Chunling Wang for viral load data, Ellen Sebastian for data processing, and Elizabeth Costello for helpful suggestions. We thank past and present members of the study team based at the Hospital Infantil Manuel de Jesús Rivera, the Centro de Salud Sócrates Flores Vivas, and the National Virology Laboratory in the Centro Nacional de Diagnóstico y Referencia of the Nicaraguan Ministry of Health, as well as the Sustainable Sciences Institute in Nicaragua for their dedication and high-quality work, and we are grateful to the study participants and their families.

This work was supported by the National Institute of Allergy and Infectious Diseases Division of Intramural Research and Division of Microbiology and Infectious Diseases (U19 AI109761, D.A.R.; U54 AI065359, A.B.), the Thomas C. and Joan M. Merigan Endowment at Stanford University (D.A.R.), the Chan Zuckerburg Biohub (D.A.R.), and by a grant (VE-1) from the Pediatric Dengue Vaccine Initiative of the Bill and Melinda Gates Foundation (E.H.). For potential conflicts of interest, see ICMJE disclosure forms.

